# Rolling the evolutionary dice: *Neisseria* commensals as proxies for elucidating the underpinnings of antibiotic resistance mechanisms and evolution in human pathogens

**DOI:** 10.1101/2023.09.26.559611

**Authors:** Kelly M. Frost, Sierra L. Charron-Smith, Terence C. Cotsonas, Daniel C. Dimartino, Rachel C. Eisenhart, Eric T. Everingham, Elle C. Holland, Kainat Imtiaz, Cory J. Kornowicz, Lydia E. Lenhard, Liz H. Lynch, Nadia P. Moore, Kavya Phadke, Makayla L. Reed, Samantha R. Smith, Liza L. Ward, Crista B. Wadsworth

**Affiliations:** Rochester Institute of Technology, Thomas H. Gosnell School of Life Sciences, Rochester, New York, USA

**Keywords:** experimental evolution, *Neisseria*, azithromycin, penicillin, antibiotic resistance, experiential learning

## Abstract

Species within the genus *Neisseria* are especially adept at sharing adaptive allelic variation across species’ boundaries, with commensal species repeatedly transferring resistance to their pathogenic relative *N. gonorrhoeae*. However, resistance in commensal *Neisseria* is infrequently characterized at both the phenotypic and genotypic levels, limiting our ability to predict novel and potentially transferable resistance mechanisms that ultimately may become important clinically. Unique evolutionary starting places of each *Neisseria* species will have distinct genomic backgrounds, which may ultimately control the fate of evolving populations in response to selection, as epistatic and additive interactions may coerce lineages along divergent evolutionary trajectories. However alternatively, similar genetic content present across species due to shared ancestry may constrain the adaptive solutions that exist. Thus, identifying the paths to resistance across commensals may aid in characterizing the *Neisseria* resistome – or the reservoir of alleles within the genus, as well as its depth. Here, we use *in vitro* evolution of four commensal species to investigate the potential for and repeatability of resistance evolution to two antimicrobials, the macrolide azithromycin and the β-lactam penicillin. After 20 days of selection, commensals evolved elevated minimum inhibitory concentrations (MICs) to penicillin and azithromycin in 11/16 and 12/16 cases respectively. Almost all cases of resistance emergence converged on mutations within ribosomal components or the *mtrRCDE* efflux pump for azithromycin-based selection, and *mtrRCDE* or *penA* for penicillin selection; thus, supporting constrained adaptive solutions despite divergent evolutionary starting points across the genus for these particular drugs. However, continuing to explore the paths to resistance across different experimental conditions and genomic backgrounds, which could shunt evolution down alternative evolutionary trajectories, will ultimately flesh out the full *Neisseria* resistome.

## INTRODUCTION

The emergence of antibiotic resistance within bacterial populations is mediated by natural selection, whereby mutations encoding drug-protective mechanisms are produced stochastically, and subsequently increase in frequency as a result of only the cells harboring these mutations surviving exposure events. However, a key question for both understanding evolutionary process and also the enhancement of surveillance efforts is: how repeatable and predictable is resistance evolution at the genotypic level? Two alternate hypotheses can be advanced: (1) adaptive landscapes are constrained to one or few solutions (i.e., genotypic constraint), or (2) a multitude of fitness peaks exist created by many mutations imparting similar phenotypic outcomes. Many prior studies support some level of genotypic constraint on resistance evolution at the strain or species-level^1–5^, however less frequently has the repeatability of resistance evolution been interrogated across species’ boundaries. Applying selection across different genomic backgrounds at the species-level may lead us to predict a higher likelihood of divergent evolutionary outcomes, with different mutations giving rise to similar phenotypic resistance in different species. We may predict this given that the pre-existing suite of potentially additive and/or epistatically-interacting mutations already present in each species’ genomes will likely be unique as a result of both genetic drift since the time of lineage divergence and also niche-specific adaptation. However, if genotypic convergence is observed across species, this suggests constrained ranges of adaptive solutions between high-level taxonomic groupings (e.g., genera, families, etc.) due to their shared ancestral history and conserved genetic makeup. Here, we begin to interrogate this question: does genotypic constraint or divergence govern the emergence of resistance evolution within the genus *Neisseria*?

The genus *Neisseria* is comprised of several Gram-negative, typically diplococcoid, oxidase-positive, and often catalase-positive species, which most frequently colonize the nasopharyngeal or oral niche in humans or animals^6^. Most human-associated *Neisseria* are carried harmlessly as commensals in 100% of healthy human adults and children, however *N. gonorrhoeae* and *N. meningitidis* are obligate and opportunistic pathogens respectively and are carried in a smaller percentage of the population (between 0.01-10%)^7–11^. Within the *N. gonorrhoeae* population, rates of resistance to multiple classes of antimicrobials are rising. For example, according to the latest Gonococcal Isolate Surveillance Project (GISP) report^12^ ∼15% of surveyed isolates were resistance to penicillin, ∼20% resistant to tetracycline, 33.2% to ciprofloxacin, 5.8% to azithromycin, and 0.3% to cefixime in the United States; and although resistance (≥ 0.25 μg/ml) was not observed in 2020 to ceftriaxone, isolates with reduced susceptibly have been identified in previous years (2017-2019) as a part of the GISP collection^12^. Additionally, surveillance studies in other countries have identified higher rates of circulating ceftriaxone resistance (e.g., 4.2% in Taiwan^13^, 16% in in Guangdong, China^14^); with recent observations indicating global dissemination (Japan, China, Europe, Australia, North America and Southeast Asia) of high-level ceftriaxone-resistant strains^15–20^. Though the genetic basis of some resistance phenotypes appears to be exclusively encoded by recurrently acquired mutations (i.e., ciprofloxacin resistance is almost always caused by amino acid substitutions in the DNA gyrase subunit A (GyrA S91F and D95G/D95A^21,22^)); the complete genetic bases of other resistance phenotypes is currently not fully described and/or is clearly imparted by multiple additive or epistatically-interacting loci (i.e., penicillin^23–27^ and azithromycin^22,28^ resistance). Thus, experimentally interrogating the paths to resistance and their repeatability will become an important component of both identifying novel contributing mutations, and understanding their potential prevalence and evolution within populations.

Studies on the paths to resistance within gonococci have previously been explored *in vitro* (e.g.,^29–34^). However, gonococci in addition to gaining resistance through *de novo* mutations, are also superbly adept at acquiring resistance from their close commensal relatives^5,28,35–37^. This allelic exchange across *Neisseria* species likely occurs in their shared colonization sites of the naso- and oropharyngeal niches^38^, with the whole genus often being referred to as a consortium with ‘fuzzy’ borders due to the high frequency of DNA donation through horizontal gene transfer (HGT)^39–41^. Commensal species thus serve as a bubbling cauldron of new adaptive solutions and reservoir of resistance for gonococci, with each species containing a unique genomic background in which novel resistance genotypes may emerge. Therefore, expanding the investigation on the repeatability of evolution to the entire genus may serve two important goals in the fight against the spread of resistance in gonococci: 1) identifying resistance phenotypes for which a multitude of genotypic paths exist, either within distinct genomic contexts or across several, and 2) determining which drugs and/or drug classes have limited adaptive solutions within the genus. Both of these findings may guide the development of nucleic acid-based resistance tests (i.e. NAAT or WGS) for surveillance programs by defining the scope of mutations which must be surveyed.

Here, we begin to interrogate the paths to resistance to two drugs with as-of-yet not fully identified genotypic bases within the pathogenic *Neisseria*. We use four different genomic contexts across the *Neisseria* genus (*N. cinerea*, *N. subflava*, *N. elongata*, and *N. canis*), and select for increasing minimum inhibitory concentrations (MICs) by passaging each species across selective gradients as previously described^5^. Though the scope of this initial and a prior study^5^ have been limited (i.e., limited species and experimental replicates) we imagine that by continuing to ‘roll the evolutionary dice’ we will ultimately coalesce on the possible and quantity of paths to resistance, addressing the repeatability of evolution to different drug classes across the genus. Finally, both this and our previous study^5^ were conducted as part of exercises within undergraduate classrooms at the Rochester Institute of Technology, highlighting the power of experimental evolution in addressing fundamental questions impacting global public health, while also providing important experiential learning opportunities for students.

## RESULTS

### Rolling the dice: Evolving *Neisseria* commensals

Four *Neisseria* commensal species were selected as distinct evolutionary starting points for antibiotic selection (*N. cinerea* (AR-0944), *N. subflava* (AR-0953 and AR-0957), *N. elongata* (AR-0945), and *N. canis* (AR-0948)). All are human-associated commensals except for *N. canis,* which colonizes the oral cavity of dogs and cats, but has also been isolated from human patients with dog and cat bite wounds^42–44^. All isolates had been phenotyped for their minimum inhibitory concentrations (MICs) to penicillin and azithromycin (Table 1), and the majority sequenced previously^45^. One isolate, AR-0944, was sequenced as a part of this study (accession: SAMN37441995; length 2.13 Mbp, 131 contigs, N50= 250 kbp, GC content 50.78%).

**Table 1.** Minimum inhibitory concetrations (MICs) for ancestral and average MICs for evolved strain.

| **Ancestral Strains** | **Azi MIC (µg/ml)** | **Average Azi**<br>**MIC (µg/ml)**<br>**evolved (n=4)** | **Pen MIC (µg/ml)** | **Average Pen**<br>**MIC (µg/ml)**<br>**evolved (n=4)** |
| --- | --- | --- | --- | --- |
| AR-0944 (*N. cinerea*) | 8 | 152 | 0.38 | 12 |
| AR-0945 (*N. elongata*) | 0.5 | 0.69 | 0.25 | 6.72 |
| AR-0948 (*N. canis*) | 0.38 | 64 | 0.25 | 5.44 |
| AR-0953 (*N. subflava*) | 2 | 224 | 1.5 † |  |
| AR-0957 (*N. subflava*) | 8 † |  | 1 | 3.69 |

† AR-0953 was only selected with azithromycin and AR-0957 was only selected with penicillin; see discussion for further details.

For each species and drug combination, four replicate lineages were passaged with selection created by application of Etest strips on standard growth media as previously described^5^ (Figure 1). Cells were passaged for 20 days, or ∼480 generations, by sweeping the entire zone of inhibition (ZOI) and a 1 cm band surrounding the ZOI, and plating any collected cells on new selective growth media. For azithromycin, the average MICs of evolved *N. cinerea* (MIC=152 ± 120.79 μg/ml), *N. canis* (64 ± 36.95 μg/ml), and *N. subflava* (224 ± 64 μg/ml) lineages crossed the breakpoint of reduced susceptibility as defined by the Clinical and Laboratory Standards Institute (CLSI) guidelines for *N. gonorrhoeae* of ≥ 2 μg/ml^46^. *N. elongata* lineages however did not surpass this breakpoint (0.69 ± 0.36 μg/ml). For penicillin, the average MICs for evolved lineages of all species surpassed the CLSI-defined breakpoint concentration of ≥ 2 μg/ml^46^: *N. cinerea* (MIC=12 ± 0 μg/ml), *N. elongata* (6.75 ± 11.53 μg/ml), *N. canis* (5.44 ± 1.38 μg/ml), and *N. subflava* (3.69 ± 2.17 μg/ml). Control populations (n=3 per species) with no drug selection showed no significant increase in azithromycin or penicillin MICs compared to the ancestral stocks (Supplementary Table 1).

**Figure 1.**
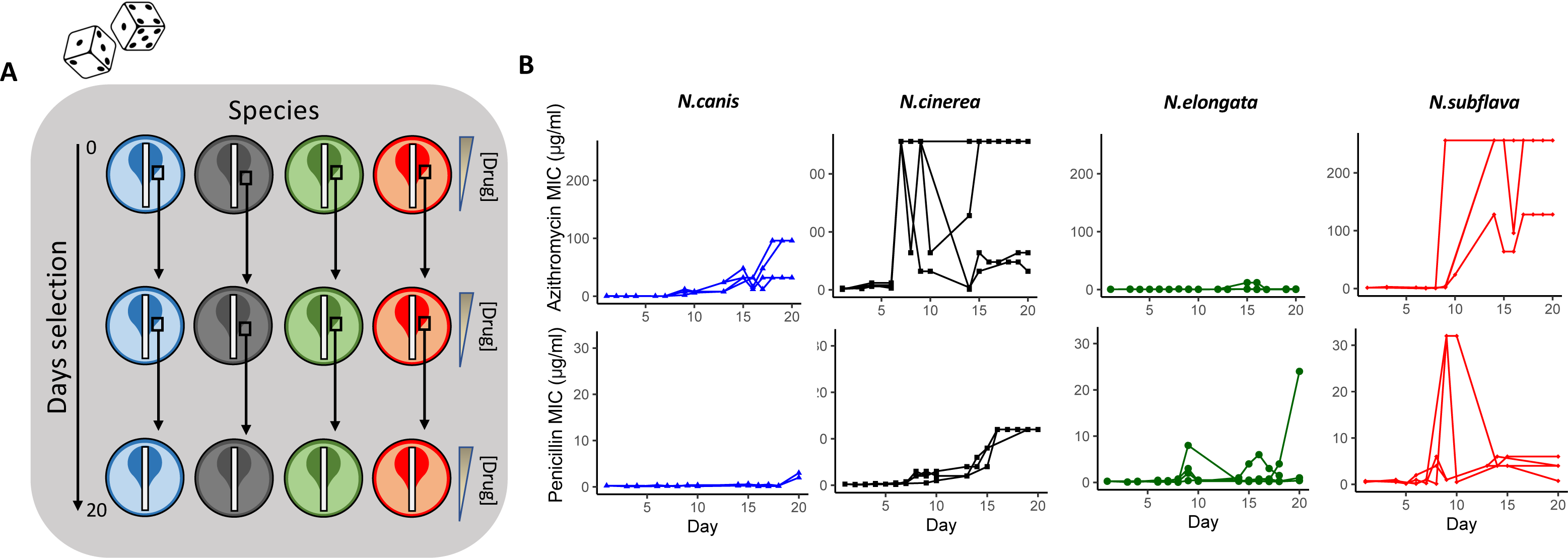
Azithromycin and penicillin-mediated selection of four species of commensal *Neisseria*. (A) Four species with distinct genetic backgrounds were selected as unique starting points for *in vitro* evolution to two antimicrobials. Each experimental replicate and species/drug combination can be envisioned as an independent “roll of the dice”, in which new derived mutations and evolutionary trajectories may emerge. In brief, 4 experimental replicates were passaged for each species and drug combination for 20 days (∼480 generations) on selective gradients created with Etest strips. Cells for each passage were selected by sweeping the entire zone of inhibition (ZOI) and a 1 cm band in the bacterial lawn surrounding the ZOI. (B) Overall, after 20 days, evolved azithromycin minimum inhibitory concentrations (MICs) tended to be higher than of penicillin MICs; with species also differing in their evolutionary trajectories towards elevated MICs within a drug class.

Final recorded MICs for azithromycin (92.17 ± 25.57 μg/ml) were significantly higher across all commensal species compared to the MICs for penicillin (4.45 ± 1.23 μg/ml) (W = 38.5, P = 0.00073; Figure 2A). Azithromycin MIC fold-changes (4.39 ± 0.77) were also significantly higher than that of penicillin MICs (2.08 ± 0.65) across species (W = 74, P = 0.043; Figure 2B). The number of days for MICs to double for azithromycin (10.75 ± 1.34) compared to penicillin (9.07 ± 0.70) were not significantly different (W = 92.5, P = 0.41; Figure 2C); nor was the day the CLSI resistance breakpoint was passed at 9.0 ± 0 and 9.0 ± 0.45 respectively (W = 18, p-value = 0.56; Figure 2D) – with species starting with above breakpoint values at the beginning of the experiment omitted for this last analysis. Between species for azithromycin, *N. subflava* and *N. cinerea* had significantly higher evolved MICs compared to *N. elongata* (Tukey’s HSD: *p* = 0.036; and *p* = 0.036 respectively; see also Figure 3A and Supplementary Table 1). There were no significant differences for final MICs between species for penicillin (Figure 3B). However, between species fold-change in MIC was significantly different for four contrasts for azithromycin (Tukey’s HSD: *p* < 0.05; Figure 3C) and three contrasts for penicillin (Tukey’s HSD: *p* < 0.01; Figure 3D).

**Figure 2.**
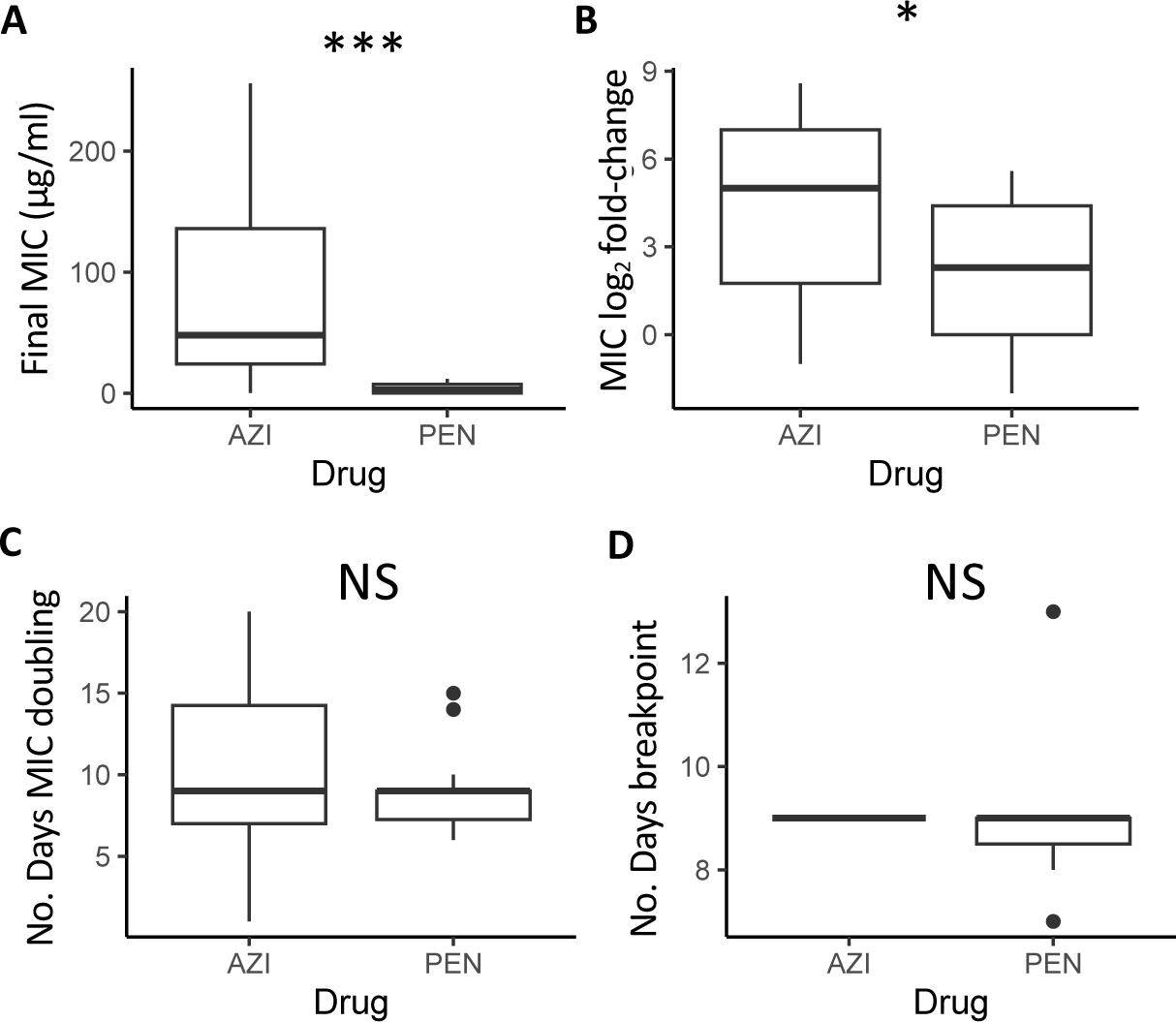
Across all species, evolved azithromycin MICs were significantly elevated compared to penicillin MICs in both (A) their final values (*p* < 0.0001), and (B) their fold-increase from ancestral MICs (*p* < 0.01). The (C) time for MICs to double was not significantly different between drugs (*p* > 0.05), as was the number of days to surpass the breakpoint value as defined by CLSI guidelines for *Neisseria gonorrhoeae* (P >0.05).

**Figure 3.**
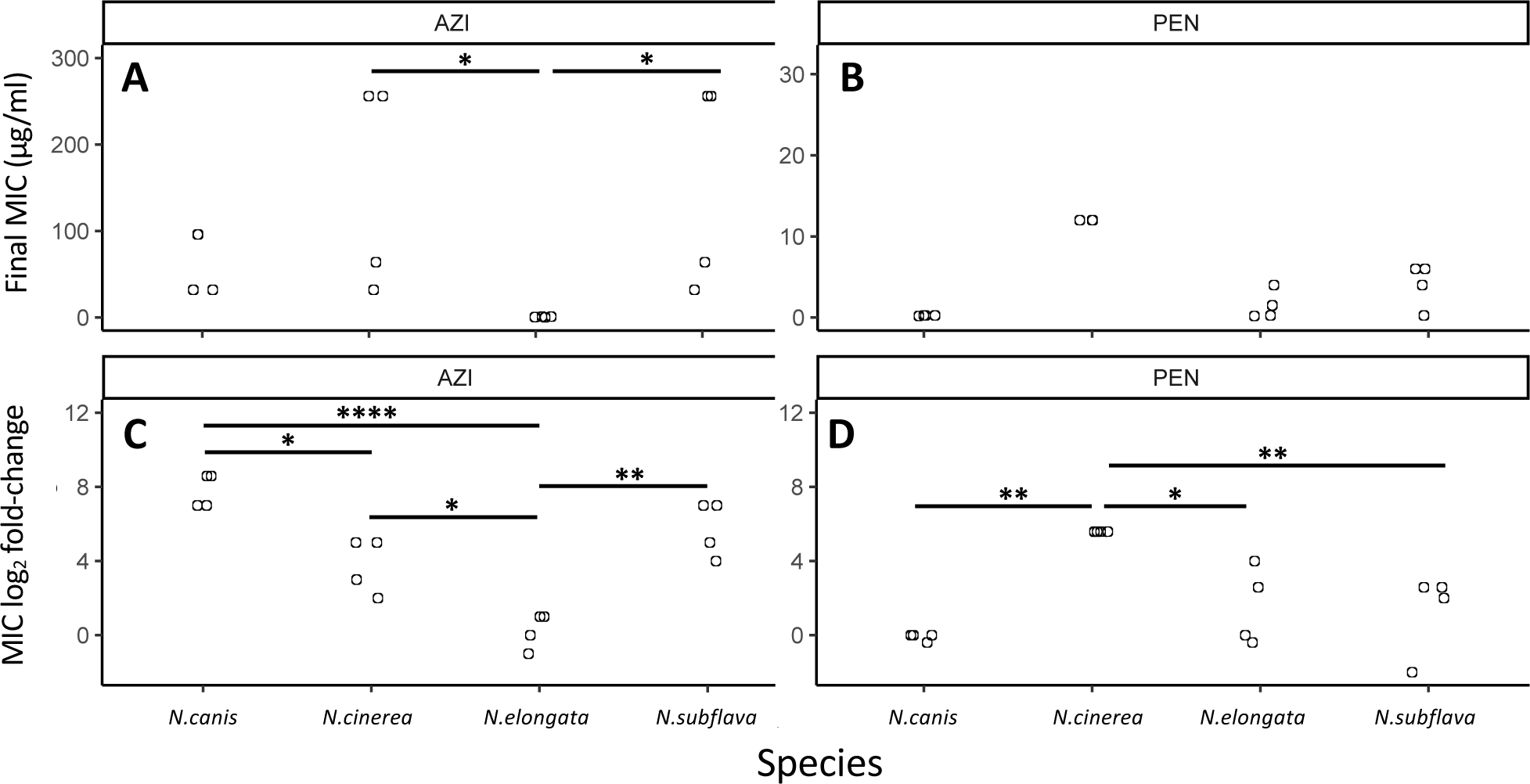
Evolved MICs and MIC log-fold change values separated by drug and species. (A) For azithromycin, *N. subflava* and *N. cinerea* had significantly higher MICs compared to *N. elongata* after selection (Tukey’s HSD: *p* = 0.036; and *p* = 0.036 respectively). (B) Species were not significantly different between any contrast for penicillin. However, between species fold-change in MIC after evolution was significantly different for (C) four contrasts for azithromycin (Tukey’s HSD: *p* < 0.05) and (D) three contrasts for penicillin (Tukey’s HSD: *p* < 0.01).

### The frequency and identity of derived mutations

For each evolved lineage, a single colony was picked for further characterization and whole-genome sequencing (Supplementary Table 1). There were no significant differences between the number of derived mutations after the 20-day long experiment between drugs across all species, however each species and interaction between drugs and species (2-way ANOVA: *p* = 0.0008) had a significant and nearly significant (2-way ANOVA: *p* = 0.055) impact on the number of derived mutations respectively. *N. elongata* had significantly fewer derived mutations compared to *N. canis* (Tukey’s HSD: *p* = 0.02), *N.* cinerea (Tukey’s HSD: *p =* 0.0007), and *N. subflava* (Tukey’s HSD: *p* = 0.004). When separated by drug class, for penicillin both *N. canis* and *N. cinerea* had significantly more derived mutations compared to *N. elongata* (Tukey’s HSD: *p =* 0.02 *and p* = 0.059 respectively; Figure 4); and for azithromycin *N. subflava* had significantly more novel mutations compared to *N. elongata* (Tukey’s HSD: *p =* 0.045; Figure 4).

**Figure 4.**
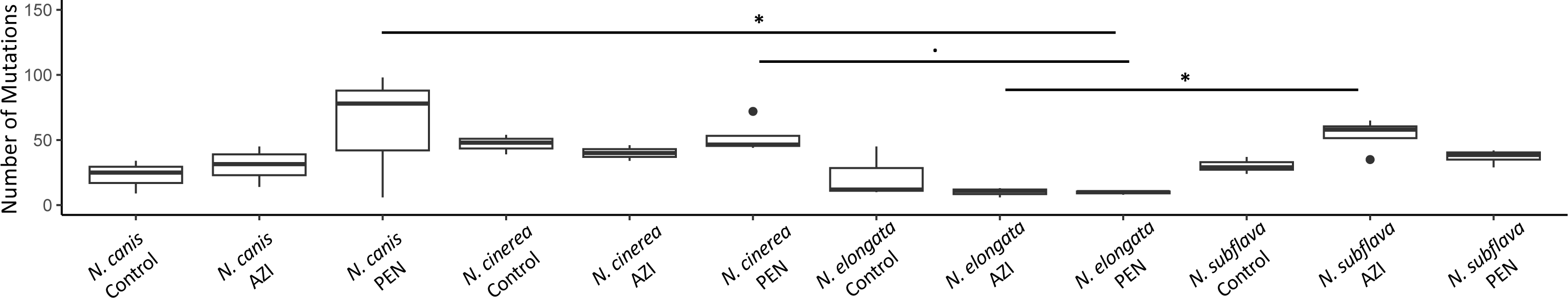
The number of derived mutations after the 20-day long experiment for azithromycin and penicillin selected lines, as well as control lineages. For penicillin both *N. canis* and *N. cinerea* had significantly more derived mutations compared to *N. elongata* (Tukey’s HSD: *p =* 0.02 *and p* = 0.059 respectively); and for azithromycin *N. subflava* had significantly more derived mutations compared to *N. elongata* (Tukey’s HSD: *p =* 0.045).

Mutations within coding domain sequences (CDSs) were identified for all evolved lineages, and after correcting for mutations also present in control lineages with no drug exposure, were considered candidates for imparting resistance (Figure 5). For azithromycin, all replicate lineages of *N. subflava*, *N. canis*, and *N. cinerea* evolved resistance, however none of the *N. elongata* strains did (Figure 1). The most frequent mutation occurring in *N. subflava* lineages was located within *pilM*. Additional mutations that emerged included those in *fabH*, *mafB5*, *nadh*, *rplP*, *rplV*, and *rpmH*. For *N.canis*, the most frequent mutations occurred in *mtrR*; followed by *rplV*, *duf2169*, *mtrD*, and *pglB2*. Finally, for *N.cinerea*, mutations emerged in *glk*, *prmA*, and *rpmH*. For penicillin, all replicate evolved lineages gained resistance except for one *N. elongata* strain and all *N.subflava* strains; however each of these lineages developed increased MICs compared to the ancestral strains, and had MICs ≥ 1 μg/ml. Mutations in *N. subflava* lineages which emerged includes those in *mtrD* and *mtrR*, *tufA*, and a *murin transglycosylase*. The most frequent mutations in *N. canis* includes those in the 16s and 23s rRNAs, followed by those in *PNL71104_P2*, *gmhA*, an HTH11-domain coding protein, a phage-associated protein, *prfB*, *rpoA*, and *tRNA-fMet(cat*). In *N. elongata*, derived mutations include those in *mtrD*, *penA*, *ispE*, and *tgt*. Finally in *N. cinerea*, mutations included those in *penA*, *pilM*, *glk*, *pitA*, *ppx*, *rpoB*, and *slmA*; along with some additional singleton mutations (Figure 5).

**Figure 5.**
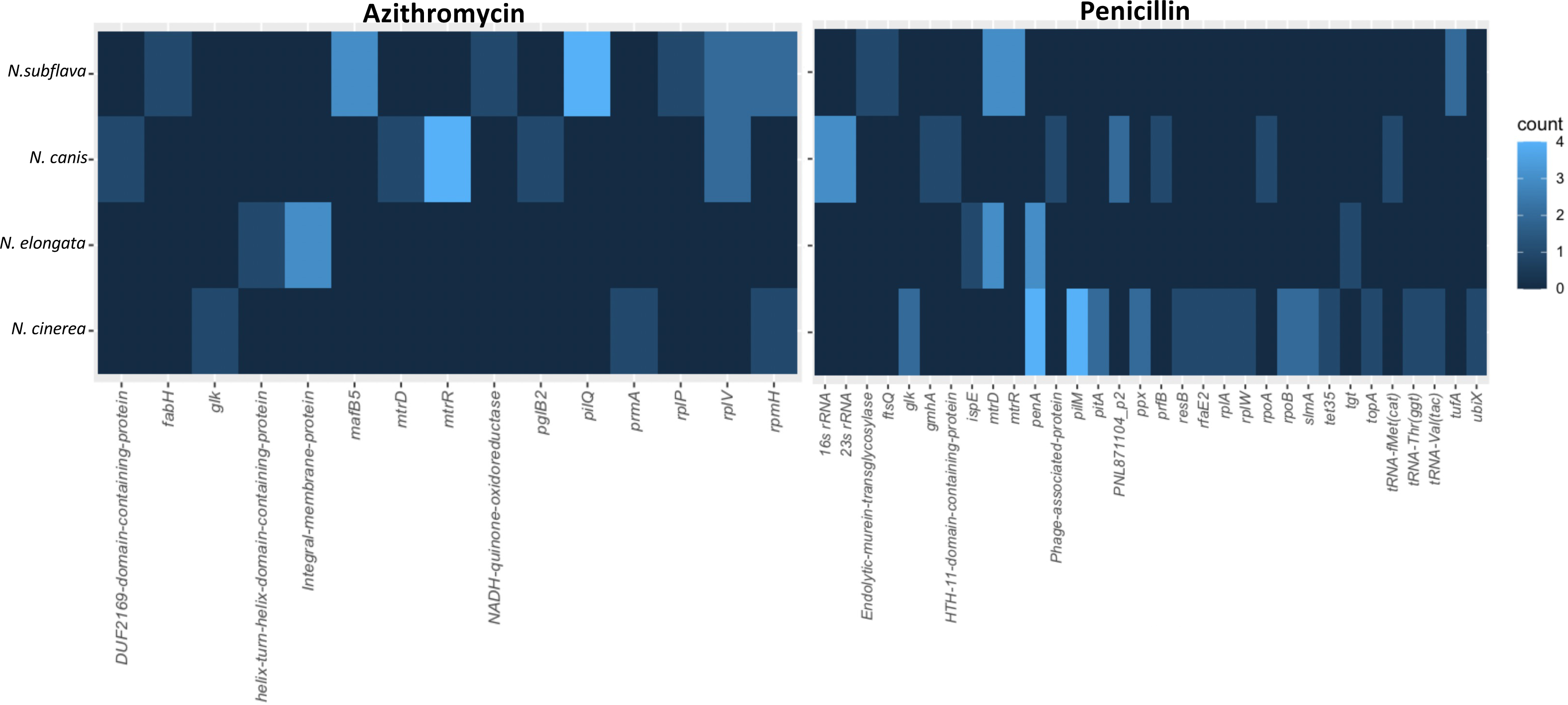
Identity of derived mutations in coding domain sequences (CDSs) for drug-selected lineages. The frequency of a mutations within a gene are displayed as a heatmap, with brighter blue coloration indicating more frequent occurrence of a mutation within a CDS in replicate evolved lineages for each species.

## DISCUSSION

Commensal *Neisseria* have repeatedly donated resistance alleles to their pathogenic relative *N. gonorrhoeae*^28,35–37^, and beyond doubt serve as a bubbling cauldron of new adaptive solutions to address ‘the antibiotic crisis’ that *N. gonorrhoeae* faces. However, we do not yet understand the full suite of resistance alleles that commensal *Neisseria* can carry, if the pool of mechanisms is large or small, and if the pool size varies by antibiotic. Here, we to role the evolutionary dice using antibiotic selection across divergent commensal *Neisseria* genomic contexts to begin to answer three important questions: 1) what are the identities of resistance mutations that can emerge in commensals, 2) are the paths to resistance evolution constrained or broad, and 3) do the answers to the two prior questions vary by drug class?

Azithromycin is a macrolide antibiotic that inhibits protein synthesis by binding to the 23S rRNA component of the 50S ribosome. Mutations that impact the conformation or block the binding site of the drug have previously been described in *N. gonorrhoeae* to impart resistance and include: mutations in the 23S rRNA azithromycin binding sites (C2611T and A2059G)^47,48^, a G70D mutation in the RplD 50S ribosomal protein L4^49^, *rplV* tandem duplications^22^, and variants of the rRNA methylase genes *ermC and ermB*^50^. Here, we also find a suite of variants that emerged post-selection within the CDSs encoding ribosomal proteins. For example, in both *N. subflava* and *N. canis* we uncovered mutations emerging in *rplV* encoding the 50S ribosomal protein L22; with 2/4 N. *subflava* lineages and 2/4 *N. canis* lineages evolving tandem duplications within this gene; previously predicted to block the azithromycin binding site^22^. In-frame insertions in *rpmH,* which encodes the 50S ribosomal L34 protein, were also frequent; and found within 2/2 surviving *N. cinerea* and 2/4 *N. subflava* strains. *N. cinerea* strains both evolved distinct *rpmH* variants (18-bp variant: GATAAGTGCGTTTCATGA; 21-bp variant: GTTGATAAGTGCGTTTCATGA), while *N. subflava* strains evolved the same variant (24-bp variant: AAACGCACTTATCAACCTTCCGTT). The *N. cinerea rpmH* variants were nearly identical to those previously described in *N. elongata*^5^ and *N. gonorrhoeae*^30^, which were found to be casual to high-level azithromycin resistance through transformation in *N. elongata*^5^, and thus are the likely mechanisms imparting high-level resistance in *N. cinerea* strains within this study. Interestingly the *N. elongata* strains evolved in this study did not evolve reduced azithromycin susceptibility (Figure 1; Table 1); however, in our prior work^5^, only 44% of replicate *N. elongata* lineages evolved resistance, and only 43% of these resistant isolates gained resistance through mutations in *rpmH.* With only 4 replicate *N. elongata* strains selected in this study we speculate that we did not have sufficient power to uncover these mutations. Finally, we find evidence for a duplication within the *rplP* gene encoding the 50S ribosomal protein L16 within a single *N. subflava* strain, however we find no difference in MICs between this strain which also harbors a *rplV* duplication and a second strain with just a *rplV* duplication, suggesting that the variant uncovered in *<u>rplP</u>* may not contribute to the elevated MICs observed. Manoharan-Basil et al. (2021)^51^ describe multiple recombination events in genes encoding ribosomal proteins across pathogenic and commensal *Neisseria*, supporting the possibility of transfer of these types of resistance mutations in natural *Neisseria* populations.

The Multiple transferable resistance efflux pump (Mtr) is a primary mechanism by which *N. gonorrhoeae* gains resistance to both azithromycin and penicillin. The Mtr efflux pump is comprised of the MtrC-MtrD-MtrE cell envelope proteins, which together export diverse hydrophobic antimicrobial agents such as antibiotics, nonionic detergents, antibacterial peptides, bile salts, and gonadal steroidal hormones from the cell^52–55^. Overexpression of the pump, through mutations that ablate or decrease the expression of the repressor of the pump (MtrR) have been demonstrated to increase resistance to both azithromycin and penicillin^22,26,56,57^; and substitutions within the inner membrane component MtrD have been shown to decrease susceptibility to azithromycin^28,36^. Here, in response to azithromycin-based selection, all four experimental replicates of *N. canis* evolved mutations in MtrR: two with a G172D substitution, one A37V, and one insertion impacting the reading frame and resulting in a premature stop codon. 3/4 replicates of *N. subflava* evolved *mtrR* mutations in response to penicillin exposure which resulted in a T11I substitution in MtrR. MtrD mutations also emerged in response to penicillin-selection in *N. subflava* (L996I) and *N. elongata* (with all three strains carrying different mutations: V139G, F604I, or A1009T). Finally, a MtrD mutation also emerged in 1/4 *N. canis* strains after azithromycin selection E823K. Interestingly, this last E823K MtrD substitution was predicted to be the causal mutation imparting azithromycin resistance in mosaic commensal Neisseria alleles transferred to *N. gonorrhoeae*^28,36^.

β-lactams, such as penicillin, target the penicillin binding proteins and inhibit cell wall biosynthesis. Mutations in Penicillin-Binding Protein 2 (PBP2, encoded by *penA*) in particular have been well documented to impart elevated penicillin MICs in *N. gonorrhoeae*^25,58^, and also other β-lactams including the extended spectrum cephalosporin ceftriaxone, through both native gonococcal alleles^59^ and non-native alleles acquired from commensal *Neisseria* ^22,37,58,60^. These mutations act by lowering the affinity of the beta-lactam antibiotics for PBP2 and also by restricting the motions of PBP2 which are important for acylation by beta-lactams^61^. Therefore, unsurprisingly we observed multiple mutations emerge in *penA*, though only in two species: *N. elongata* and *N. cinerea*. 3/3 surviving *N. elongata* evolved lines had *penA* mutations emerge: P399S, V574E, and A581S; and all four experimental *N. cinerea* replicates evolved *penA* mutations encoding the amino acid substitutions: F518S, V548E, and A549E.

Additional derived mutations of note that emerged after selection include those in the RNA polymerase and components of the pilus. Here, after penicillin selection a *rpoA* mutation emerged in *N. canis,* and *rpoB* mutations emerged in *N. cinerea*. In *N. gonorrhoeae*, both RpoD (E98K and Δ92) and RpoB (R201H) mutations impact ceftriaxone susceptibility, likely through increased expression of PBP1 and reduced expression of D,D-carboxypeptidase^62^. Here, the *rpoA* G147A nucleotide substitution in *N. canis* resulted in a silent change so does not likely contribute to elevated penicillin MICs; however, the evolved *rpoB* mutations did encode amino acid substitutions (E345A and P591S) in 2/4 *N. cinerea* replicate lineages. Finally, the pilus-associated mutations in PilM in *N. cinerea* in response to penicillin selection and PilQ in *N. subflava* in response to azithromycin likely impact drug diffusion across the outer membrane in some way similar to gonococci^63^, however are not likely to be evolutionarily maintained in natural *Neisseria* populations due to the importance of the pilus in host-cell attachment^64^.

The aforementioned ribosomal, MtrRCDE, and PenA mechanisms seem to be the likely contributors to the emergence of reduced susceptibly in all of the *Neisseria* commensals investigated in this study for both penicillin and azithromycin-based selection (Figure 6). Therefore, despite 2/2 *N. canis* replicates evolving low-level penicillin resistance with as-of-yet unexplained genetic bases; with 19/21 cases of *Neisseria* evolution converging on known resistance mechanisms, we must accept a constrained range of adaptive solutions to antibiotic selection within the genus at this point. Remaining questions do exist however. For example: MICs varied greatly among experimental replicates of the same species, so what other modulating mutations emerged that impact resistance phenotype? Furthermore, here we only investigate coding-domain regions, thus important mutations in intergenic regions were likely missed (i.e., promoter region mutations). We also acknowledge that our small sample of strains and experimental replicates may have limited the pool of potential resistance mechanisms uncovered. For example, some mechanisms may be less frequently observed due to high fitness costs, necessitating the evolution of compensatory mutations. These types of mutations may therefore be missed in small-scale experimental studies. Finally, evolution does not occur in controlled laboratory environments, so what is the role of intergenus gene exchange in *Neisseria* resistance emergence? Can other genera transfer clinically relevant resistance mechanisms to the *Neisseria* (see Goytia & Wadsworth (2022)^35^ for a discussion on this possibility)? In summary, our current results highlight conserved paths to resistance within the *Neisseria* genus, though continued tosses of the evolutionary dice may ultimately paint a different picture.

**Figure 6.** Paths to resistance emergence across members of the *Neisseria* genus. For azithromycin selection, all species with evolved resistance converged on mutations within ribosome components or the *mtrRCDE* efflux pump system. For penicillin resistance, *N.cinerea*, *N. elongata*, and *N. subflava* all strains evolving resistance acquired mutations in either the *mtrRCDE* efflux pump system or *penA*. *N. canis* experimental replicates evolving penicillin resistance acquired as-of-yet undescribed resistance mutations.

## METHODS

### Bacterial strains and culturing

Stocks of *Neisseria* were obtained from the Centers for Disease Control and Prevention (CDC) and Food and Drug Association’s (FDA) Antibiotic Resistance (AR) Isolate Bank “*Neisseria* species MALDI-TOF Verification panel”. Evolved strains included: AR-0944 (*N. cinerea*), AR-0945 (*N. elongata*), AR-0948 (*N. canis*), AR-0953 (*N. subflava*), and AR-0957 (*N. subflava*). Bacteria were cultivated for all subsequent protocols on GC agar base (Becton Dickinson Co., Franklin Lakes, NJ, USA) media plates containing 1% Kelloggs solution (GCP-K plates) for 18-24 hours at 37°C in a 5% CO_2_ atmosphere. Bacterial stocks were stored in trypticase soy broth (TSB) containing 50% glycerol at −80°C.

### Experimental evolution and MIC testing

Minimum inhibitory concentrations (MICs) were measured by Etest strips (bioMérieux, Durham, NC) on GCB-K plates according to the manufacturer specifications. In brief, cells from overnight plates were suspended in TSB to a 0.5 McFarland standard and inoculated onto GCB-K plates. Etest strips were incubated on these plates for 18-24 hours at 37°C in a 5% CO_2_ incubator. MICs were subsequently determined by reading the lowest concentration that inhibited growth of bacterial lawns.

For each of the four *Neisseria sp.* used in the study, four replicates were passaged on GCB-K plates containing a selective gradient of either penicillin or azithromycin. Selective gradients were created using Etest strips as described above and previously^5^, and MICs were recorded each day. Cells to be passaged were collected from the entire zone of inhibition (ZOI) and a 1 cm region in the bacterial lawn surrounding the ZOI (Figure 1). Cells were suspended in TSB, and spread onto a new GCB-K plate containing a fresh Etest strip. Strains were exposed to azithromycin and penicillin for 20 days, or ∼480 generations. Controls for each species were passaged on GCB-K plates as described above, however they did not contain any antibiotic.

### Genomic sequencing and comparative genomics

DNA was isolated from cells using the PureLink Genomic DNA Mini kit (Thermo Fisher Corp., Waltham, MA), following lysis in TE buffer (10 mM Tris [pH 8.0], 10 mM EDTA) with 0.5 mg/mL lysozyme and 3 mg/mL proteinase K (Sigma-Aldrich Corp., St. Louis, MO). Resultant genomic DNA was treated with RNase A and prepared for sequencing using the Nextera XT kit (Illumina Corp., San Diego, CA). Libraries were uniquely dual-indexed and pooled, and sequenced on the Illumina MiSeq platform at the Rochester Institute of Technology Genomics Core using V3 600 cycle cartridges (2×300bp). Sequencing quality of each paired-end read library was assessed using FastQC v0.11.9^65^. Trimmomatic v0.39^66^ was used to trim adapter sequences, and remove bases with phred quality score < 15 over a 4 bp sliding window. Reads < 36 bp long, or those missing a mate, were also removed from subsequent analysis. Draft assemblies had been previously published for all strains^45^, except for *N. cinerea* AR-0944. This assembly was constructed using SPAdes v.3.14.1^67^ and all assemblies were annotated with Bakta v.1.8.1^68^. Assembly quality was assessed using QUAST (http://cab.cc.spbu.ru/quast/). Trimmed reads were mapped back to draft assemblies using Bowtie2 v.2.2.4^69^ using the “end-to-end” and “very-sensitive” options and Pilon v.1.16^70^ was used to call variant sites. Data analysis and visualizations were conducted in R^71^.

### Data Availability

All scripts and datasets are available on: https://github.com/wadsworthlab. Read libraries for the genomics datasets generated in this study can be accessed on the Sequence Read Archive for evolved strains can be access as a part of the BioProject PRJNA1018855. The assembly for AR-0944 has been deposited to GenBank (accession: SAMN37441995).

## Acknowledgements

This work was produced by the members of the Fall 2022 Genomics course (BIOL340) at the Rochester Institute of Technology (RIT) – a big thank you to every student for their effort in the design and implementation of these experiments. The authors would like to acknowledge the generous support provided by the RIT College of Science and the Thomas H. Gosnell School of Life Science for this work. Work reported in this publication was also supported by the National Institute of General Medical Sciences of the National Institutes of Health under Award Number R15GM149587. The content is solely the responsibility of the authors and does not necessarily represent the official views of the National Institutes of Health. The funders had no role in study design, data collection and analysis, decision to publish, or preparation of the manuscript. The authors would also like to thank Girish Kumar at the RIT Genomics Core for providing support and sequencing services.

## Supplementary Figure Captions

**Supplementary Figure 1**. Ancestral azithromycin MICs started significantly higher across species compared to penicillin MICs (P < 0.001).

