## Supplemental Table 1 for "Rolling the evolutionary dice: *Neisseria* commensals as proxies for elucidating the underpinnings of antibiotic resistance mechanisms and evolution in human pathogens"

**Supplemental Table 2. The impact of species on evolved MICs, 2-way ANOVA interaction results ordered by adjusted P-value**

| Comparision | Adjusted P |
|---|---|
| #AZI:N.elongata-AZI:N.cinerea | 0.03599 |
| #AZI:N.subflava-AZI:N.elongata | 0.03599 |
| #AZI:N.cinerea-AZI:N.canis | 0.4938929 |
| #AZI:N.subflava-AZI:N.canis | 0.4938929 |
| #AZI:N.elongata-AZI:N.canis | 0.8257135 |
| #PEN:N.cinerea-PEN:N.canis | 0.9999931 |
| #PEN:N.elongata-PEN:N.cinerea | 0.9999968 |
| #PEN:N.subflava-PEN:N.cinerea | 0.9999995 |
| #PEN:N.elongata-PEN:N.canis | 1 |
| #PEN:N.subflava-PEN:N.canis | 1 |
| #AZI:N.subflava-AZI:N.cinerea | 1 |
| #PEN:N.subflava-PEN:N.elongata | 1 |

**Supplemental Table 3. The impact of species on MIC fold-change, 2-way ANOVA interaction results ordered by adjusted P-value**

| Comparision | Adjusted P |
|---|---|
| #AZI:N.elongata-AZI:N.canis | 0.0000018 |
| #PEN:N.cinerea-PEN:N.canis | 0.0001473 |
| #AZI:N.subflava-AZI:N.elongata | 0.0002319 |
| #PEN:N.subflava-PEN:N.cinerea | 0.0045885 |
| #AZI:N.cinerea-AZI:N.canis | 0.0084122 |
| #PEN:N.elongata-PEN:N.cinerea | 0.0085094 |
| #AZI:N.elongata-AZI:N.cinerea | 0.0300176 |
| #AZI:N.subflava-AZI:N.canis | 0.4613159 |
| #AZI:N.subflava-AZI:N.cinerea | 0.4870153 |
| #PEN:N.elongata-PEN:N.canis | 0.7070757 |
| #PEN:N.subflava-PEN:N.canis | 0.8444889 |
| #PEN:N.subflava-PEN:N.elongata | 0.9999949 |
