## Supplemental Table 2 and 3 for "Rolling the evolutionary dice: *Neisseria* commensals as proxies for elucidating the underpinnings of antibiotic resistance mechanisms and evolution in human pathogens"

**Supplementary Table 1. MICs and number of derived mutations compared to ancestral reference sequences for single colony picks of evolved lineages**

| Species | Strain | Selective agent | MIC of derived strains after selection |
|---|---|---|---|
| *N. canis* | AR0948.C1 | Azi | 0.38 |
| *N. canis* | AR0948.C2 | Azi | 0.38 |
| *N. canis* | AR0948.C3 | Azi | 0.38 |
| *N. canis* | AR0948A.1 | Azi | 16 |
| *N. canis* | AR0948B.1 | Azi | 48 |
| *N. canis* | AR0948C.1 | Azi | 48 |
| *N. canis* | AR0948D.1 | Azi | 32 |
| *N. canis* | AR0948.C1 | Pen | 0.25 |
| *N. canis* | AR0948.C2 | Pen | 0.25 |
| *N. canis* | AR0948.C3 | Pen | 0.25 |
| *N. canis* | G4.S1.1 | Pen | 3 |
| *N. canis* | G4.S2.1 | Pen | † |
| *N. canis* | G4.S3.2 | Pen | 2 |
| *N. canis* | G4.S4.1 | Pen | † |
| *N. cinerea* | AR0944.C1 | Azi | 8 |
| *N. cinerea* | AR0944.C2 | Azi | 6 |
| *N. cinerea* | AR0944.C3 | Azi | 8 |
| *N. cinerea* | G2.S1.1 | Azi | † |
| *N. cinerea* | G2.S2.1 | Azi | 256 |
| *N. cinerea* | G2.S3.1 | Azi | † |
| *N. cinerea* | G2.S4.1 | Azi | 256 |
| *N. cinerea* | AR0944.C1 | Pen | 0.38 |
| *N. cinerea* | AR0944.C2 | Pen | 0.38 |
| *N. cinerea* | AR0944.C3 | Pen | 0.38 |
| *N. cinerea* | AR0944A.1 | Pen | 12 |
| *N. cinerea* | AR0944B.1 | Pen | 4 |
| *N. cinerea* | AR0944C.1 | Pen | 2 |
| *N. cinerea* | AR0944D.1 | Pen | 6 |
| *N. elongata* | AR0945.C1 | Azi | 0.5 |
| *N. elongata* | AR0945.C2 | Azi | 0.75 |
| *N. elongata* | AR0945.C3 | Azi | 0.5 |
| *N. elongata* | AR0945A.1 | Azi | † |
| *N. elongata* | AR0945B.1 | Azi | 1 |
| *N. elongata* | AR0945C.1 | Azi | 0.5 |
| *N. elongata* | AR0945D.1 | Azi | 0.25 |
| *N. elongata* | AR0945.C1 | Pen | 0.25 |
| *N. elongata* | AR0945.C2 | Pen | 0.25 |
| *N. elongata* | AR0945.C3 | Pen | 0.25 |
| *N. elongata* | G5.S1.1 | Pen | 1 |
| *N. elongata* | G5.S2.1 | Pen | † |
| *N. elongata* | G5.S3.1 | Pen | 2 |
| *N. elongata* | G5.S4.1 | Pen | 12 |
| *N. subflava* | G1.S1.1 | Azi | 256 |
| *N. subflava* | G1.S2.1 | Azi | 256 |
| *N. subflava* | G1.S3.1 | Azi | 96 |
| *N. subflava* | G1.S4.1 | Azi | 96 |
| *N. subflava* | G3.S1.1 | Pen | 4 |
| *N. subflava* | G3.S2.1 | Pen | 6 |
| *N. subflava* | G3.S3.1 | Pen | 4 |
| *N. subflava* | G3.S4.1 | Pen | 0.75 |
| *N.subflava* | AR0953.C1 | Azi | 2 |
| *N.subflava* | AR0953.C2 | Azi | 1 |
| *N.subflava* | AR0953.C3 | Azi | 1 |
| *N.subflava* | AR0957.C1 | Azi | 8 |
| *N.subflava* | AR0957.C2 | Azi | 8 |
| *N.subflava* | AR0957.C3 | Azi | 8 |
| *N.subflava* | AR0953.C1 | Pen | 1 |
| *N.subflava* | AR0953.C2 | Pen | 1.5 |
| *N.subflava* | AR0953.C3 | Pen | 1.5 |
| *N.subflava* | AR0957.C1 | Pen | 1 |
| *N.subflava* | AR0957.C2 | Pen | 1 |
| *N.subflava* | AR0957.C3 | Pen | 1 |

† Stocks of single colony picks did not grow after the initial patch plate
